# Establishment of a CPER Reverse Genetics System for Powassan Virus Defines Attenuating NS1 Glycosylation Sites and an Infectious NS1-GFP11 Reporter Virus

**DOI:** 10.1101/2023.05.03.539311

**Authors:** Jonas N. Conde, Grace E. Himmler, Megan C. Mladinich, Yin Xiang Setoh, Alberto A. Amarilla, William R. Schutt, Nicolas Saladino, Elena E. Gorbunova, Daniel J. Salamango, Eckard Wimmer, Hwan Keun Kim, Erich R. Mackow

## Abstract

Powassan virus (POWV) is an emerging tick-borne Flavivirus that causes lethal encephalitis and long term neurologic damage. Currently there are no POWV therapeutics, licensed vaccines or reverse genetics systems for producing infectious POWVs from recombinant DNA. Here we used a circular polymerase extension reaction (CPER) approach to generate recombinant LI9 (recLI9) POWVs with attenuating NS1 protein mutations and a recLI9-split-eGFP reporter virus. Flavivirus NS1 proteins are highly conserved glycoproteins that regulate replication, spread and neurovirulence. POWV NS1 proteins contain three putative N-linked glycosylation sites that we modified individually in infectious recLI9 mutants (N85Q, N208Q, N224Q). NS1 glycosylation site mutations reduced replication kinetics and were attenuated with a 1-2 log decrease in infectious titers. The severely attenuated recLI9-N224Q mutant exhibited a 2-3 day delay in focal cell-to-cell spread and reduced NS1 secretion. Like WT LI9, the recLI9-N224Q mutant was lethal when intracranially inoculated into suckling mice. However, footpad inoculation of recLI9-N224Q resulted in the survival of 80% of mice and demonstrated that NS1-N224Q mutations attenuate POWV neuroinvasion *in vivo*. To monitor NS1 trafficking, we CPER fused a split GFP11-tag to the NS1 C-terminus and generated an infectious reporter virus, recLI9-NS1-GFP11. Cells infected with recLI9-NS1-GFP11 revealed NS1 trafficking in live cells and the novel formation of large NS1 lined intracellular vesicles. An infectious recLI9-NS1-GFP11 reporter virus permits real-time analysis of NS1 functions in POWV replication, assembly and secretion, and provides a platform for evaluating antiviral compounds. Collectively, our robust POWV reverse genetics system permits analysis of viral spread and neurovirulence determinants *in vitro* and *in vivo*, and enables the rational genetic design of live attenuated POWV vaccines.

## Significance

Our findings newly establish a mechanism for genetically modifying POWVs, systematically defining pathogenic determinants and rationally designing live attenuated POWV vaccines. This initial study demonstrates that mutating POWV NS1 glycosylation sites attenuates POWV spread and neurovirulence *in vitro* and *in vivo.* Our findings validate a robust circular polymerase extension reaction (CPER) approach as a mechanism for developing, and evaluating, attenuated genetically modified POWVs. We further designed an infectious GFP-tagged reporter POWV that permits us to monitor secretory trafficking of POWV in live cells, and which can be applied to screen potential POWV replication inhibitors. This robust system for modifying POWVs permits provides the ability to define attenuating POWV mutations and create genetically attenuated recPOWV vaccines.

## Introduction

Flaviviruses (FVs) are enveloped, positive-strand RNA viruses that are globally distributed, primarily transmitted by arthropod vectors (mosquitoes and ticks) and infect an estimated 400 million people annually(^1, 2^). Mosquito-borne FVs cause an array of symptoms ranging from mild fevers, rashes and jaundice to acute hemorrhagic fever, shock, microcephaly and encephalitic syndromes(^2^). While FVs are widely distributed by mosquito vectors, tick-borne FVs are spread in discrete geographic locations and constrained by the range of tick vectors and their sustaining mammalian hosts(^1^). Tick-borne FVs cause encephalitic human diseases, with Tick-borne encephalitis virus (TBEV) responsible for an estimated 10-15,000 encephalitis cases per year in Eurasia. Powassan virus (POWV) is the only tick-borne FV in North America(^1^) and is emerging as a cause of encephalitis in the U.S.(^1–3^). Currently, there are no approved vaccines or therapeutics for preventing or resolving POWV neurovirulence, and live attenuated recombinant POWV vaccines have yet to be developed(^1, 2^).

POWV is a neurovirulent tick-borne flavivirus present in tick saliva and is transmitted by injection into tick bite sites in as little as 15 minutes(^1, 4^). In symptomatic patients, POWV causes severe encephalitis with a 10% mortality rate, and 50% of cases have debilitating long-term neurologic damage(^1^). POWV, strain LB, was initially isolated from the brain of a 5 year old in Powassan, Ontario Canada in 1958(^5, 10, 74^), and since then POWVs have been found widely in *Ixodes* ticks(^3, 6–16^). Two POWV genetic lineages are spread by discrete *Ixodes* ticks, but comprise a single POWV serotype with 96% identical envelope proteins providing common vaccination targets(^1, 2, 13, 14, 16^). Determinants of POWV spread and neurovirulence remain to be determined, but are constrained by the lack of POWV isolates derived directly from ticks, without passage in murine brains, and reverse genetics approaches for generating and comparing infectious recombinant POWV mutants.

The emergence of POWV disease parallels the spread of tick vectors in North America(^2, 3, 6, 12, 17^), and recent analysis revealed ∼2% POWV prevalence in Long Island, NY deer ticks(^18^). POWV strain LI9 was isolated from Long Island deer ticks by directly infecting VeroE6 cells, without murine neuroadaptation(^5, 10, 13, 19^). In epithelial cells, POWV LI9 nonlytically spreads cell-to-cell forming infected cell foci, and in mice LI9 is neurovirulent and elicits cross-reactive neutralizing POWV antibody responses(^19^). How POWV spreads to neuronal compartments remains to be revealed, but the ability of POWVs to hemagglutinate(^10^) provides a conduit for hematogenous dissemination. *In vitro*, POWVs infect primary human brain microvascular endothelial cells (hBMECs) and, without permeabilizing polarized monolayers, POWVs are basolaterally released from hBMECs(^19^). Basolateral release from hBMECs provides a potential mechanism for POWVs to spread across the blood-brain-barrier (BBB) into protected neuronal compartments.

Flavivirus RNAs (∼11 kb) encode a single open reading frame that is cotranslationally processed by viral and cellular proteases into 3 structural proteins (Capsid, prM and Envelope) and seven nonstructural (NS) proteins(^2, 20, 21^). FVs replicate in the cytoplasm, bud into the lumen of the ER and are released from cells in exocytic vesicles. Envelope (Env) and NS1 proteins are both translocated into the lumen of the ER, yet only integral membrane glycoprotein Env is assembled onto the virion(^31, 35, 38^). Env proteins direct cellular attachment and viral entry, and are the primary targets of protective neutralizing antibodies(^2, 21–25^). In contrast to Env, translocated NS1 is glycosylated and secreted from cells, and required for cytoplasmic FV replication(^26–37^). Env and NS1 proteins both contain conserved and FV specific glycosylation sites, and changes in the glycosylation of either protein can impact FV virulence(^31, 35, 38^).

NS1 is translocated into the ER lumen where it is glycosylated, dimerizes, and secreted from cells as a lipophilic hexameric complex(^26, 27, 39, 40^). Within the ER, NS1 peripherally associates with, and remodels, lumenal ER membranes and nucleates the assembly of cytoplasmic FV polymerase complexes through interactions with integral membrane NS2A/4A/4B proteins(^27, 28, 30, 40–42^). NS1 conservation across FV groups reflects multifunctional roles for NS1 domains in ER membrane curvature, viral replication, NS1 secretion, dimer and hexamer assembly, lipid binding, and many suggested pathogenic functions: binding complement, cell permeability and viremia(^27, 28, 30, 31, 33, 35, 36^, ^37, 41–44^). The array of proposed NS1 functions highlights the importance of NS1 in pathogenesis, and rationalizes mutating NS1 as a mechanism of FV and POWV attenuation(^31, 35, 38, 41, 45–47^).

Analysis of Japanese encephalitis virus (JEV) NS1 mutants distinguished NS1 residues required for replication (AA160) from viral particle formation (AA273)(^36^). However, mutating specific NS1 N-linked glycosylation sites (NxS/T) has been reported to either enhance or inhibit the replication and neurovirulence of discrete FVs(^34, 45, 46, 48^). All FV NS1 proteins have an N-linked glycosylation site at residue 207/208(^26, 31, 34, 38, 39, 49^), however, POWV NS1 proteins also contain two novel glycosylation sites (N85 and N224)(^31^). Functions of unique POWV NS1 glycosylation sites have yet to be evaluated, but may play key roles in POWV replication, secretion, cell-to-cell spread and pathogenesis.

Reverse genetics systems are needed to genetically modify POWVs, define determinants of POWV pathogenesis and create live attenuated recombinant POWV vaccines. Typical reverse genetics approaches are based on cloning viral RNA genomes into bacterial plasmids or artificial chromosome vectors with the toxicity and stability issues of large viral inserts that magnify the complexity of genetic virus modification (^50–56^). The circular polymerase extension reaction (CPER) was originally devised to clone complex genes(^57, 58^) and subsequently used to bypass the plasmid instability of large viral inserts and to rapidly generate mutant and reporter viruses(^36, 59–64^). CPER has now been used to investigate virulence determinants in West Nile virus (WNV), Zika virus (ZIKV), Yellow Fever and Dengue virus (DENV) and JEV(^36, 59–63, 65, 66^), and has the potential to be applied to other positive stranded RNA viruses(^61, 67^).

Here we used CPER to develop a robust POWV reverse genetics system and generate an infectious LI9 POWV from cDNA (recLI9) that mirrors WT LI9 in sequence, replication and focal cell-to-cell spread in VeroE6 cells. CPER was used to generate recLI9 mutant viruses lacking individual NS1 glycosylation sites (N85Q; N208Q; N224Q), and to generate an infectious recLI9 split-Green fluorescent protein (GFP) reporter virus(^68–72^). In contrast to LI9, mutating LI9 NS1 glycosylation sites reduced replication kinetics, viral titers and delayed focal cell-to-cell spread *in vitro*. The recLI9-NS1-N224Q mutant was severely attenuated with foci formation delayed by 2-3 days *in vitro,* and in vivo resulted in the survival of 80% of peripherally inoculated mice. These findings define a key role for the NS1-N224 glycosylation site in POWV replication, cell-to-cell spread and pathogenesis; and suggest N224Q as an attenuating mutation to be considered in the development of live attenuated POWV vaccines.

In order to monitor NS1 functions during POWV infection, we used CPER to add a split-GFP tag to the NS1 C-terminus and generate an infectious, GFP expressing, recLI9 reporter virus(^68–72^). Infecting cells expressing ER-GFP1-10 cells with the POWV GFP11 reporter virus (recLI9-NS1-GFP11) reconstituted GFP fluorescence and permitted real-time live cell analysis of NS1 spread during POWV infection. Our findings revealed perinuclear NS1 ER trafficking in POWV infected cells, as well as the novel accumulation of NS1-GFP in large intracellular vesicles. In the context of focal POWV spread, NS1 accumulation in vesicles suggests a potential mechanism for POWV NS1 to uniquely regulate viral release and cell-to-cell spread(^68, 73^). In addition to NS1 trafficking, an infectious recLI9-NS1-GFP11 reporter virus provides a fluorescence-based assay for screening potential inhibitors of POWV replication and maturation(^68, 69, 71, 72^). Collectively, a robust POWV reverse genetics system permits us to systematically define determinants of POWV pathogenesis, evaluate attenuating POWV mutations and create genetically modified recPOWVs for use as live attenuated vaccines.

## Results

### POWV Spreads Cell-to-Cell in the Presence of Neutralizing Antibody

The direct isolation of POWV LI9 in VeroE6 cells revealed a novel focal spread phenotype that occurred in the absence of restrictive overlays and without lytic plaque formation(^19^). POWVs bud into the lumen of the ER and are trafficked via secretory vesicles to the cell surface where they remain largely cell associated(^10, 74^). Foci formation suggested the potential for POWV to spread cell-to-cell similar to Hepatitis C virus (HCV), a distant FV family member, that spreads focally cell-to-cell in a neutralizing antibody independent manner(^75–77^). To investigate whether LI9 foci are formed by cell-to-cell spread, we evaluated foci formation in the presence of POWV neutralizing antibody (ATCC-HMAF) or control antibody added daily to infected cell supernatants. Anti-POWV antibody added to cells during viral adsorption prevented infection, while the post-adsorption addition of control or anti-POWV antibody at a 1:250 dilution (neutralizing titer 1:6400) failed to inhibit the formation of POWV infected cell foci (Fig. 1A). This establishes that POWV LI9 initially spreads cell-to-cell and explains the focal spread of LI9 during isolation from ticks, and during passage in epithelial cells in liquid culture. These are novel findings for tick-borne FVs that may permit POWV to bypass immune surveillance(^78^), and provide potential mechanisms for POWVs to spread at tick bite sites or into the CNS.

**Figure 1.**
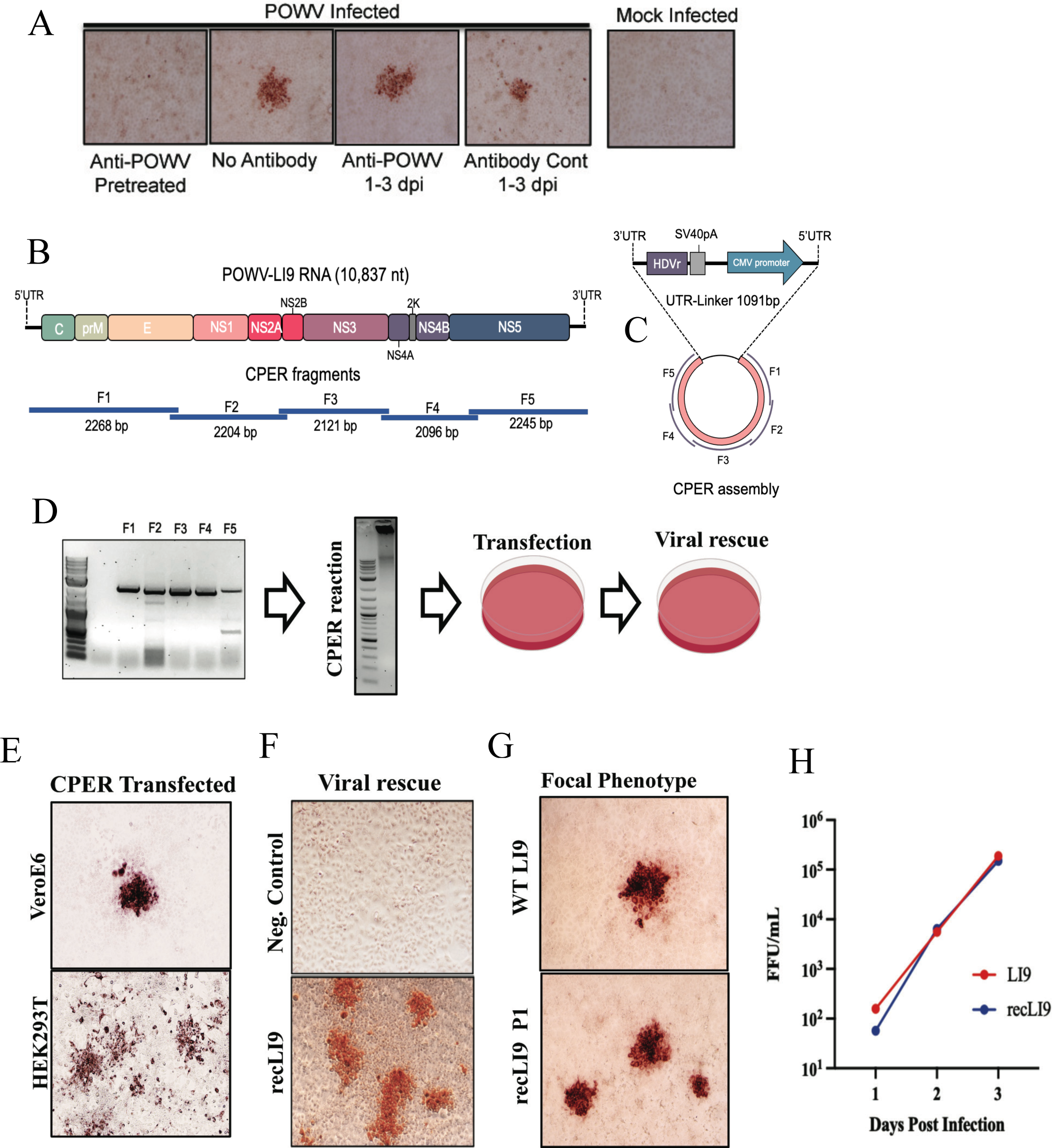
CPER Generation and Cell-to-Cell Spread of Recombinant POWV Strain LI9. **(A)** VeroE6 cells were infected with LI9 POWV at an MOI of 0.01 with or without anti-POWV HMAF (1:250) during adsorption. Cells were washed with PBS and media was replaced with DMEM with or without anti-POWV HMAF (1:250), or control ascitic fluid 6 hours post adsorption. Cells were methanol fixed 3 dpi and immunostained with anti-POWV HMAF (1:5,000)(^19, 97, 98^). **(B)** Schematic of the LI9 POWV genome and overlapping fragments amplified from LI9 cDNA. **(C)** CPER assembly schematic of F1-F5 fragments with a UTR-Linker fragment containing the last 26 nucleotides of the LI9 3’UTR, hepatitis delta virus ribozyme (HDVr), SV40 polyadenylation signal, a CMV promoter and 33 nucleotides of the LI9 5’UTR. **(D)** Agarose gel electrophoresis of PCR-amplified fragments (F1-F5) showing a representative image of three experimental repeats. F1-F5 were combined in equal molar amounts with the UTR-Linker in a CPER reaction. A representative agarose gel of CPER reaction product. Resultant LI9 CPER products were transfected into HEK293T or VeroE6 cells and supernatants were subsequently used to infect VeroE6 cells and rescue infectious recLI9 viruses. **(E)** Immunostaining CPER transfected VeroE6 or HEK293T cells (7 or 3 dpt, respectively) display focal cell-to-cell spread foci morphologies. **(F)** Infectious recLI9 virus rescued from CPER transfected cells and grown in HEK293T cells, versus CPER controls amplified without the UTR-linker fragment. **(G)** Comparison of WT LI9 and recLI9 focal cell-to-cell spread phenotypes in immunostained VeroE6 cells. **(H)** Growth kinetics of WT LI9 (red) and recLI9 (blue) POWVs 1-3 dpi in VeroE6 cells (MOI 1).

### CPER Reverse Genetics – A Mechanism for Defining Determinants of POWV Pathogenesis

Currently, there are no established mechanisms for attenuating POWVs or defining determinants of POWV neurovirulence. To generate recombinant POWVs (revPOWVs), we used a CPER approach(^36, 59–64^) to develop a robust POWV reverse genetics system (Fig. 1B-D). POWV LI9 RNA was initially used to generate first-strand cDNA that was amplified into five fragments comprising the full-length POWV genome and a sixth circularizing UTR-linker fragment (Fig 1B). Amplified fragments contain 26 nucleotide overlapping sequences, with the UTR-linker fragment containing 26 nt from the LI9 3’UTR, a hepatitis delta virus ribozyme (HDVr), SV40 polyadenylation signal, CMV promoter, and the first 33 nt of the 5’UTR sequence (Fig. 1C). To generate circularized CPER DNAs, LI9 F1-F5 cDNA fragments and the UTR-linker DNA were amplified using high-fidelity Phusion DNA polymerase (Fig 1D). CPER amplified DNA was purified and directly transfected into HEK293T or VeroE6 cells. Three days post-transfection, supernatants from transfected cells were inoculated into VeroE6 cells and infectious viral rescue was evaluated by immunoperoxidase staining of LI9 infected cells (Fig 1E, F). Cells infected by recLI9 retained a focal cell-to-cell spread phenotype, replication kinetics and titers, with sequences identical to WT LI9 (Fig. 1G). These results demonstrate the authenticity of the CPER generated recLI9 POWV and the development of a robust reverse genetic system for generating POWVs from recombinant DNA.

### CPER Mutagenesis of Specific NS1 Glycosylation Sites Inhibits POWV Spread and Replication

NS1 is translocated into the lumen of the ER where it both orchestrates cytoplasmic replication complex assembly and ER-Golgi virion maturation(^31, 35, 36^). NS1 is modified by N-linked glycosylation with high-mannose carbohydrates that direct dimerization(^31, 35, 38, 39^), and Golgi compartment processing of complex oligosaccharides facilitates NS1 secretion(^39, 79^). NS1 proteins from mosquito-borne FVs contain two conserved N-linked glycosylation sites (N130 and N207/208) (^26, 31, 38, 80^). In contrast, tick-borne FV NS1 proteins have three N-linked glycosylation sites (N85, N207/208, N223/224)(Fig. 2A,B) with 2 novel putative sites absent from other FVs (^31, 81^). The location of NxT sites on NS1 dimers is presented on putative POWV LI9 NS1 protein structures predicted using AlphaFold2 on the LI9 NS1 (Fig 2B,S2).

**Figure 2.**
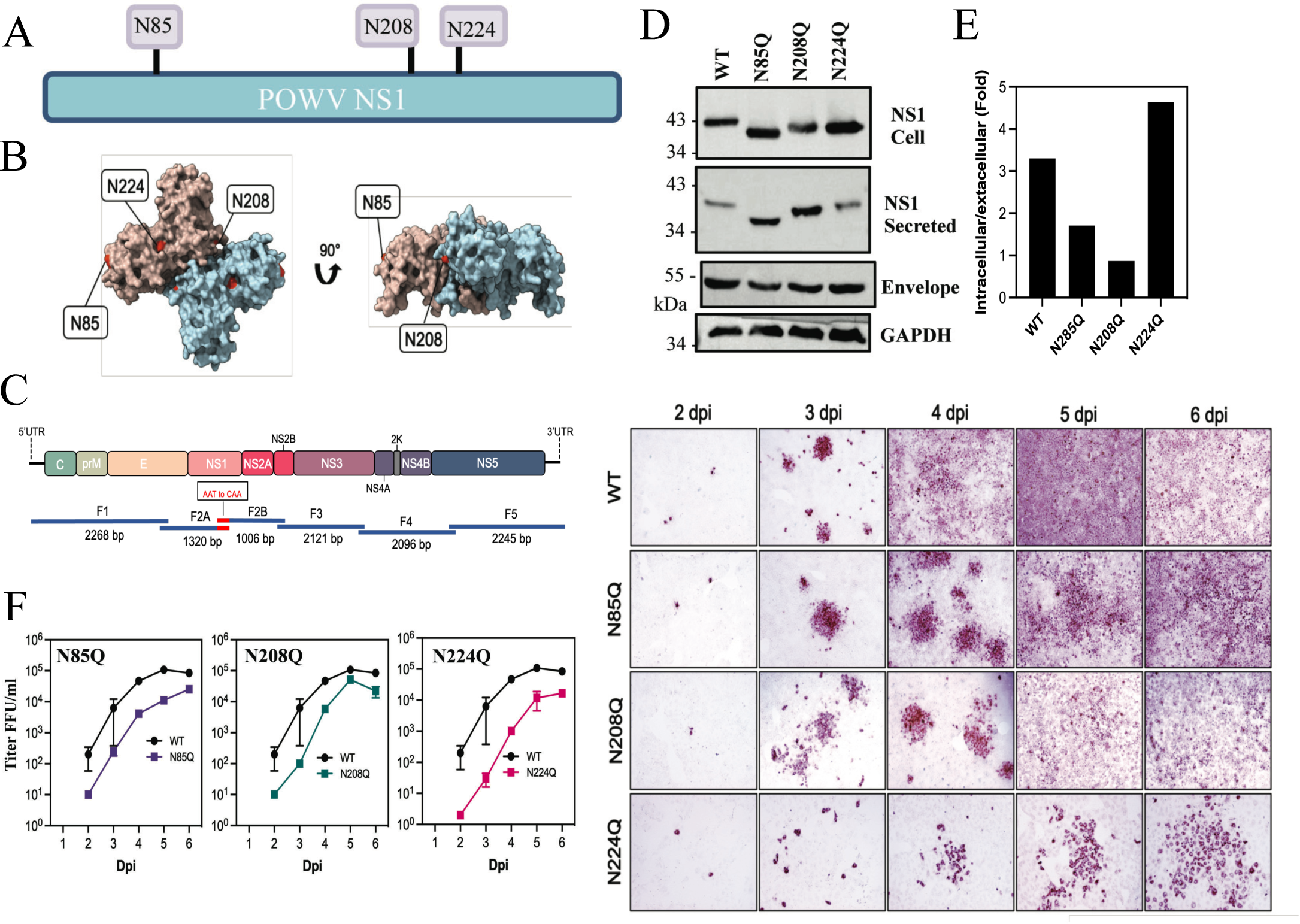
CPER POWV NS1 glycosylation mutants have impaired replication kinetics. **(A)** Schematic of N-linked glycosylation sites within POWV NS1 proteins. **(B)** Alphafold2 model of dimeric POWV NS1 protein showing pink or blue monomers and the localization of putative N-linked glycosylation sites (N85, N208, N224) in red^(100)^. **(C)** Schematic strategy for CPER generating N-linked glycosylation site (NxT) mutants using overlapping fragments to introduce N to Q codon changes. Fragment 2 was split into subfragments 2A and 2B with overlapping regions incorporating mutations. **(D)** VeroE6 cells were infected with WT LI9, recLI9-NS1_N85Q_, recLI9-NS1_N208Q_, and recLI9-NS1_N224Q_ mutant viruses (MOI 1) and 7 dpi cell lysates, and supernatants, were analyzed by Western blot using antibodies to POWV-NS1 (mAb), anti-POWV HMAF or GAPDH. **(E)** Ratios of intracellular versus extracellular NS1 levels from rec-LI9-NS1 glycosylation mutants. **(F)** Growth kinetics of WT LI9 (black) and CPER-generated NS1 glycosylation mutants recLI9-NS1_N85Q_ (purple), recLI9-NS1_N208Q_ (green), and recLI9-NS1_N224Q_ (pink), 1-6 dpi of VeroE6 cells (MOI 0.1). **(G)** Kinetic comparison of focal cell-to-cell spread by WT LI9 and recLI9-NS1 glycosylation mutants 1-6 dpi in VeroE6 cells (MOI 0.1) by anti-POWV (HMAF) immunoperoxidase staining of POWV infected cells.

Here we used POWV reverse genetics to modify individual POWV NxS/T sites (N85Q, N208Q, or N224Q) to determine whether NS1 glycosylation affects viral replication and focal cell-to-cell spread. Using CPER, we split F2, containing NS1, into F2A and F2B using overlapping primers that contain glycosylation site specific N->Q mutations (Fig 2C). Amplified F2A and F2B fragments were substituted for F2 in CPER reactions and used to generate recLI9-NS1 NxT mutants, as in Figure 1. Sequencing of rescued recLI9-NS1 mutants verified site-specific N->Q residue changes within NS1 proteins of each individual POWV (N85Q, N208Q and N224Q) (Fig. S1). To investigate roles for NS1 NxT sites in NS1 expression and secretion, we infected VeroE6 cells at an MOI of 10 with the LI9-NS1-N85Q;-N208Q; or - N224Q mutant viruses. Intracellular and secreted NS1 proteins were evaluated 6 dpi by immunoblotting using an anti-NS1 mouse monoclonal antibody (Figure 2D,E). Compared to WT LI9, mutating N208Q and N224Q glycosylation sites increased migration of NS1 proteins by Western Blot, while migration of NS1 from the N85Q mutant was dramatically increased from both intracellular and secreted NS1 sources (Figure 2D). Evaluation of the relative amounts of intracellular and secreted NS1 proteins revealed a 200-300% increase in the secretion of N85Q and N208Q NS1 proteins and a ∼50% decrease in the secretion of the N224Q NS1 protein compared to NS1 from WT LI9 (Fig 2E).

Relative to WT LI9 infection, titers of recLI9 NS1 glycosylation mutants revealed slower viral replication kinetics with maximal titers reduced by 1-3 logs from 2-6 dpi (Fig. 2F). While each NS1 glycosylation mutant spread focally, the formation of foci in monolayers was delayed by 1 day in N85Q and N208Q NS1 mutants (Fig 2G). In contrast, foci formation was reduced by 2-3 days in the LI9-NS1-N224Q mutant, which failed to spread uniformly throughout monolayers by 6 dpi (Fig 2G). These findings indicate that mutating any single LI9 NS1 glycosylation site reduced viral replication and spread, while mutating the NS1-N224Q glycosylation site resulted in a highly attenuated recLI9 mutant that was severely restricted in replication and spread *in vitro*.

Attempts to CPER generate recPOWV LI9 mutants with two glycosylation site changes (N85Q/N208Q; N85Q/N224Q; or N208Q/N224Q) failed to result in recovery of any double mutants. It remains unclear whether additional glycosylation site mutations critically impact NS1 functions required for POWV replication and virion assembly or whether double mutants could be rescued by transcomplementation in NS1 protein expressing cells.

### POWV NS1 Mutant Dimerization, Glycosylation and Secretion

Changing NS1 protein glycosylation may impact dimer formation, glycan processing and NS1 protein functions. To determine whether dimer formation is altered in POWV NS1 glycosylation mutants, we compared untreated or heat treated (2 min at 100°C) NS1 dimer stability prior to SDS-PAGE analysis. We found that all mutants formed dimers in unheated samples and that both dimers and monomers of glycosylation mutants were formed, but with increased migration on SDS gels versus WT LI9 NS1 (Figure 3A). These findings are consistent with unaltered dimer formation by DENV NS1 glycosylation site mutants(^38^).

**Figure 3.**
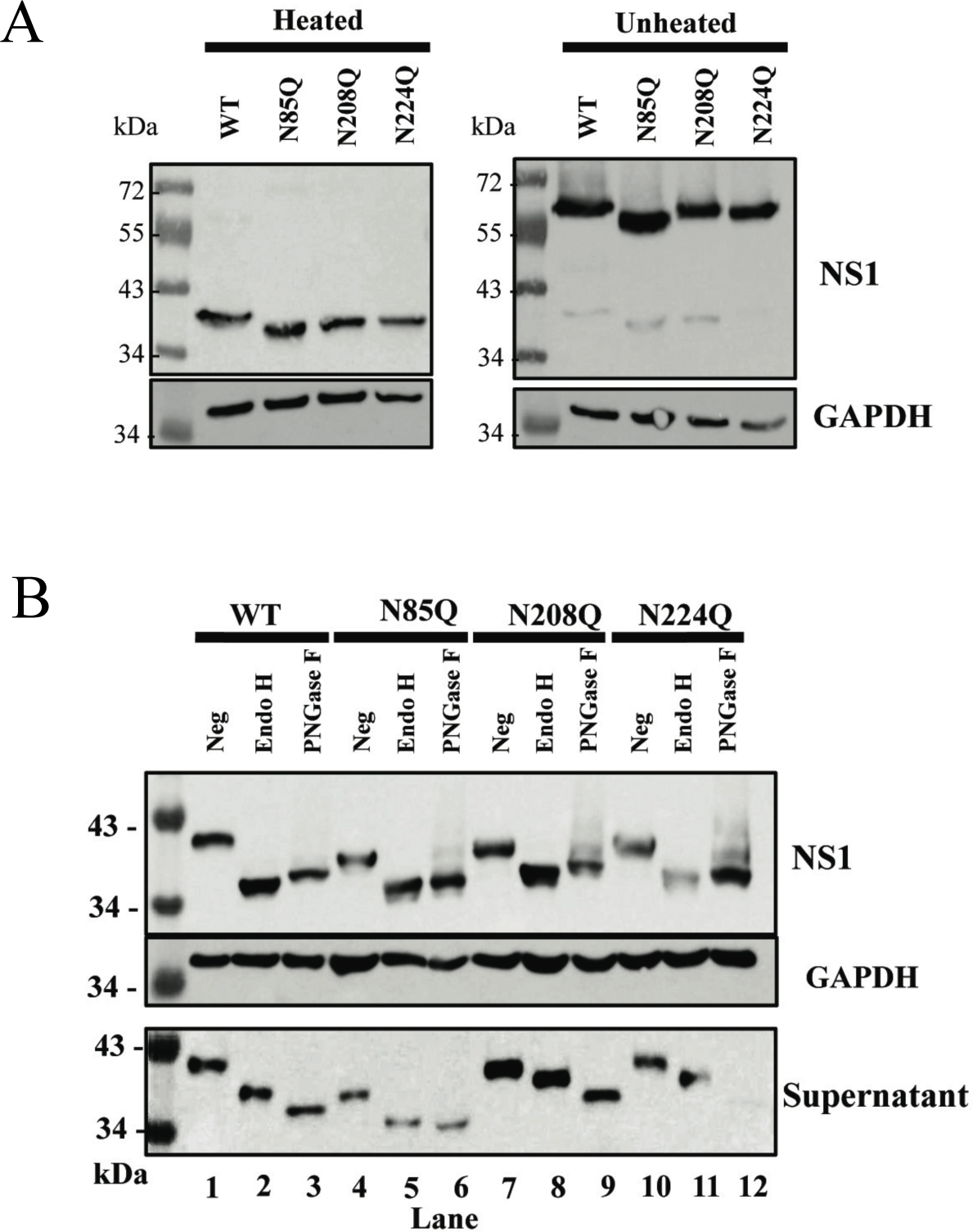
POWV NS1 Mutant Dimerization and Glycosidase Analysis. **(A)** VeroE6 cells were infected with WT POWV LI9, recLI9-NS1_N85Q_, recLI9-NS1_N208Q_, or recLI9-NS1_N224Q_ mutants (MOI, 1). Cell lysates were harvested 7 dpi with or without heat directed dimer disassembly prior to Western blot analysis using anti-POWV-NS1 mAb (1:5,000), or anti-GAPDH (1:5,000). **(B)** VeroE6 cells were infected with WT LI9, recLI9-NS1_N85Q_, recLI9-NS1_N208Q_, or recLI9-NS1_N224Q_ mutant viruses (MOI, 1). Cell lysates were harvested 7 dpi and 20 µg of protein lysate or 50 µL of supernatant was subjected to Endo H or PNGase F digestion. Undigested and digested samples from cell lysate or supernatant were analyzed by Western blot using antibody to POWV-NS1 or GAPDH.

We evaluated changes in high mannose or complex glycan addition to N-linked glycans of LI9 NS1 glycosylation mutants using endoglycosidases. Cell-associated and secreted NS1 proteins from WT and NxT mutants were treated with, Endo H to remove high-mannose N-linked glycans, or PNGase F to remove a combination of high mannose, hybrid, and complex oligosaccharides. Digestion profiles demonstrate that intracellular NS1 from WT LI9 or recLI9 glycosylation mutants are largely composed of high-mannose N-linked glycans as digestion with either Endo H or PNGase F resulted in similar NS1 protein migration (Fig. 3B). However, secreted NS1 proteins of WT, N208Q and N224Q are only partially reduced in size following Endo H digestion compared to PNGase (Fig. 3B) suggesting that N208Q and N224Q mutant NS1 glycans are further modified in the secretory pathway to contain complex or hybrid sugars (Fig. 3B). In contrast, the secreted NS1 N85Q protein migrates similarly after Endo H or PNGase F digestion indicating that glycans on NS1 N85Q mutant proteins are not further processed in the Golgi (Figure 3B). The large size difference of N85Q mutants, compared to NS1 proteins from LI9 WT or N208Q, and N224Q mutants, suggests that the N85 glycosylation site determines whether N208 and N224 glycans are further processed during NS1 secretion.

### Analysis of Attenuated recLI9-NS1-N224Q in Mice

POWV LI9 is lethal when intracranially or subcutaneously inoculated into C57BL/6 mice(^19^). Here we assessed the lethality of *in vitro* attenuated recLI9-NS1-N224Q in murine models and found that like WT LI9, recLI9-NS1-N224Q is 100% lethal when i.c. inoculated into suckling mice (N=5) 5-7 dpi. Although both WT LI9 and recLI9-N224Q are lethal when i.c. inoculated, there was a 1 day difference in the timing of lethality between WT and mutant (Fig. 4A,B). As i.c. inoculation of suckling mice reflects neurovirulence, but not neuroinvasion, we s.c. inoculated C57BL/6 mice with LI9 or recLI9-NS1-N224Q (N=8-10) and evaluated weight loss and lethality from 1-20 dpi. WT LI9 caused weight loss and was lethal in 40% of C57BL/6 mice 9-12 dpi. In contrast, in mice s.c. inoculated with recLI9-NS1-N224Q lethal disease onset was delayed, with 80% of mice surviving infection (Fig. 4B,C). Similar to attenuating replication and spread kinetics *in vitro*, our findings suggest that mutating the N224Q glycosylation site in infectious LI9 is attenuating *in vivo*, and partially reduces lethal POWV outcome.

**Figure 4.**
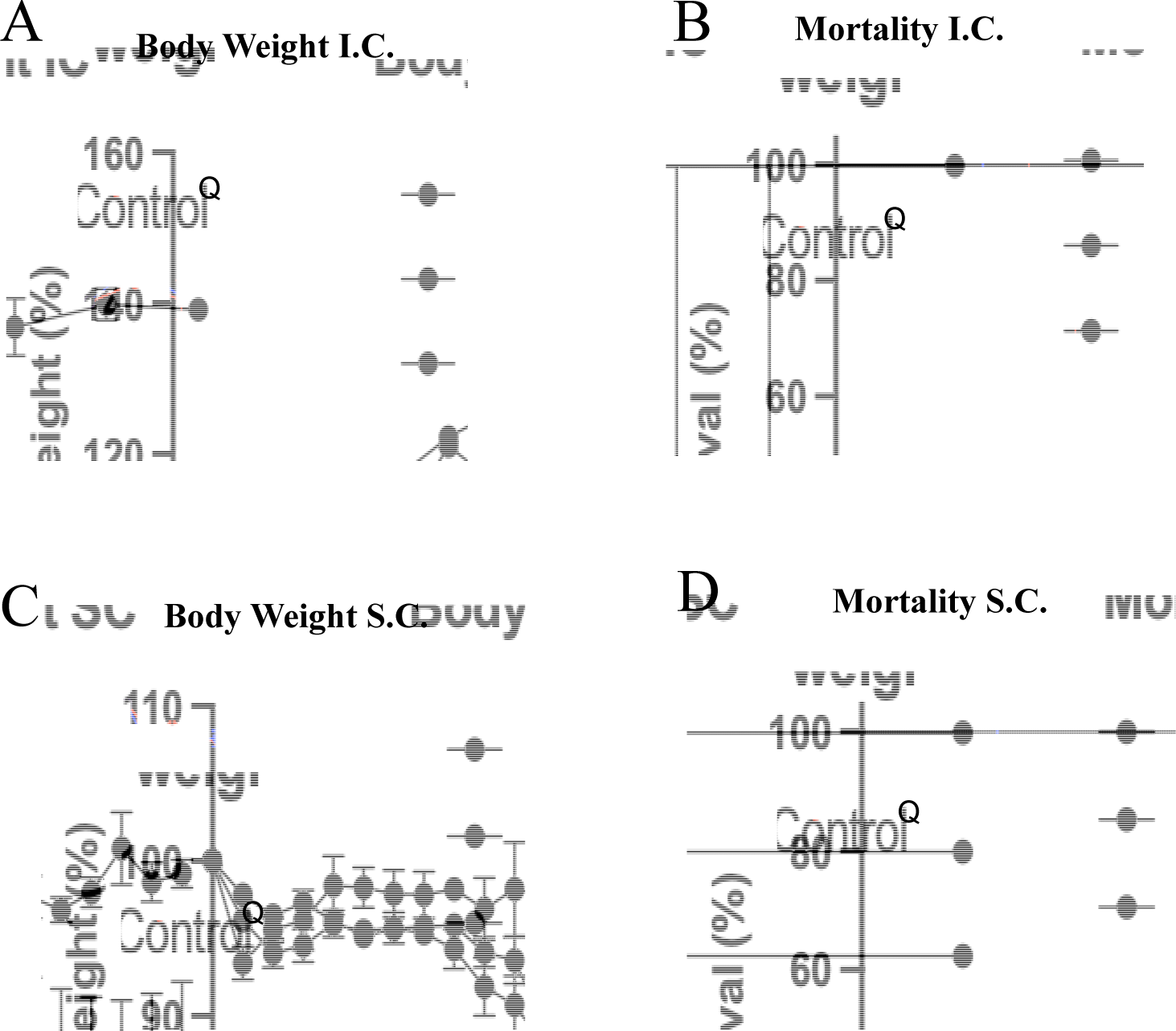
POWV N224Q Mutation Attenuates Neurovirulence in C57BL/6 mice. **(A)** Body weight analysis and **(B)** Kaplan-Meier survival curves (LI9 vs recLI9-N224Q, P=0.0442) of pups of C57BL/6 mice (N=3-5/ group) intracranially inoculated with 2 × 102 FFU of WT LI9, recLI9-N224Q, or buffer-only. **(C)** Body weight analysis and **(D)** Kaplan-Meier survival curves (LI9 vs. K224Q, P=0.3310) for C57BL/6 mice (male, N=4-10/group) footpad inoculated with 2 × 103 FFU of WT LI9, recLI9-N224Q or a buffer-only control.

### Generation of an Infectious POWV Split11-GFP Reporter Virus

Recombinant viruses expressing fluorescent proteins permits live cell, real time, analysis of cellular protein interactions, viral protein trafficking and the development of high throughput screens for viral inhibitors (^63, 65, 68, 70, 72, 73, 82, 83^). Expressing the 11^th^ β-strand of green fluorescent protein (GFP11-residues 215-230) with coexpressed GFP_1-10_ (residues 1-214) reconstitutes a fluorescent GFP molecule (Fig. 5A)(^68, 69, 71^). As a means of analyzing POWV NS1 protein functions, we used CPER to generate NS1 C-terminally tagged with a split GFP11 fusion protein in recLI9 (recLI9-GFP11) (Fig. 5B). The recLI9-GFP11 reporter virus forms infected cell foci by immunostaining and replicates with similar kinetics and titers 1-3 dpi as WT LI9 in VeroE6 cells (Fig. 5C,D). HEK293T and VeroE6 cells were transduced with retroviruses expressing cytoplasmic GFP_1-10_ and mCherry (mCh-GFP_1-10_), or an ER translocated-GFP_1-10_-KDEL and cytoplasmic mCherry (mCh-ER-GFP_1-10_). We infected mCh-GFP_1-10_ or mCh-ER-GFP_1-10_ expressing HEK293T cells with the recLI9-NS1-GFP11 virus and found GFP positive infected cell foci in ER-GFP_1-10_ but not in GFP_1-10_ expressing cells 2-3 dpi (Fig 5E,S3). Similarly, in VeroE6 cells expressing mCh-ER-GFP_1-10,_ infection with recLI9-NS1-GFP11 reconstituted NS1-GFP fluorescence with a perinuclear localization consistent with NS1-GFP11 translocation to the ER (Fig. 5F). In addition, we found abundant expression of NS1-GFP11 in large fluorescent vesicles (6 dpi) that may reflect NS1 protein accumulation in intracellular secretory or specialized vesicles. Intracellular NS1 accumulation provides a rationale for reduced NS1 secretion during POWV infection and suggests that novel vesicular trafficking may foster cell-to-cell spread in POWV infected cells. Collectively, using CPER reverse genetics, we have generated an infectious recLI9-GFP11 reporter virus that provides a mechanism for analyzing the role of NS1 in restricting POWV secretion or directing cell-to-cell spread, and provides an infectious fluorescent reporter virus that can be used to screen POWV inhibitors.

## Discussion

We isolated the LI9 strain of POWV directly from ticks in VeroE6 cells in order to avoid potential neuroadaptive changes of prototype POWV strains (LB and SP) that were isolated and adapted to growth in murine brains(^5, 10, 13, 19^). The LI9 POWV isolate nonlytically infects VeroE6 cells, unexpectedly spreading focally in cells lacking restrictive overlays(^19^). LI9 spread focally in media containing high levels of anti-POWV neutralizing antibodies, revealing that LI9 spreads cell-to-cell without requiring apical release into cell supernatants (Fig. 1). This both confirms the largely cell associated nature of POWVs(^10^), while uniquely mirroring data establishing that HCV, a distantly related FV, spreads cell-to-cell in hepatocytes(^75-78^). The neutralizing antibody independent, cell-to-cell, spread of HCV is proposed to occur at partially sealed cell junctions that permit HCV to evade immune surveillance(^75–78^). Measles virus (MV) was also shown to spread cell-to-cell in epithelial cells, without syncytia formation, using a GFP reporter virus(^84^). The MV-GFP reporter was visualized flowing to the cytoplasm of adjacent cells through lateral pores and was suggested as a mechanism for MV transmission from epithelial cells to neurons(^78, 84^). Implying that POWV cell-to-cell spread is epithelial cell specific, we found that POWV fails to spread cell-to-cell in hBMECs or pericytes. At later times post-infection, POWV titers increase and spreads conventionally by the diffusion of cell-free viral particles(^78^). Cell-to-cell spread is a known viral mechanism for immune evasion(^78^) that may contribute to dissemination in the skin or foster CNS transmission via epithelial cell barriers that comprise the choroid plexus(^85–87^). The mechanism of POWV cell-to-cell spread remains to be defined, and our recLI9-GFP reporter virus may permit analysis of POWV trafficking and spread to recipient cells.

**Figure 5.**
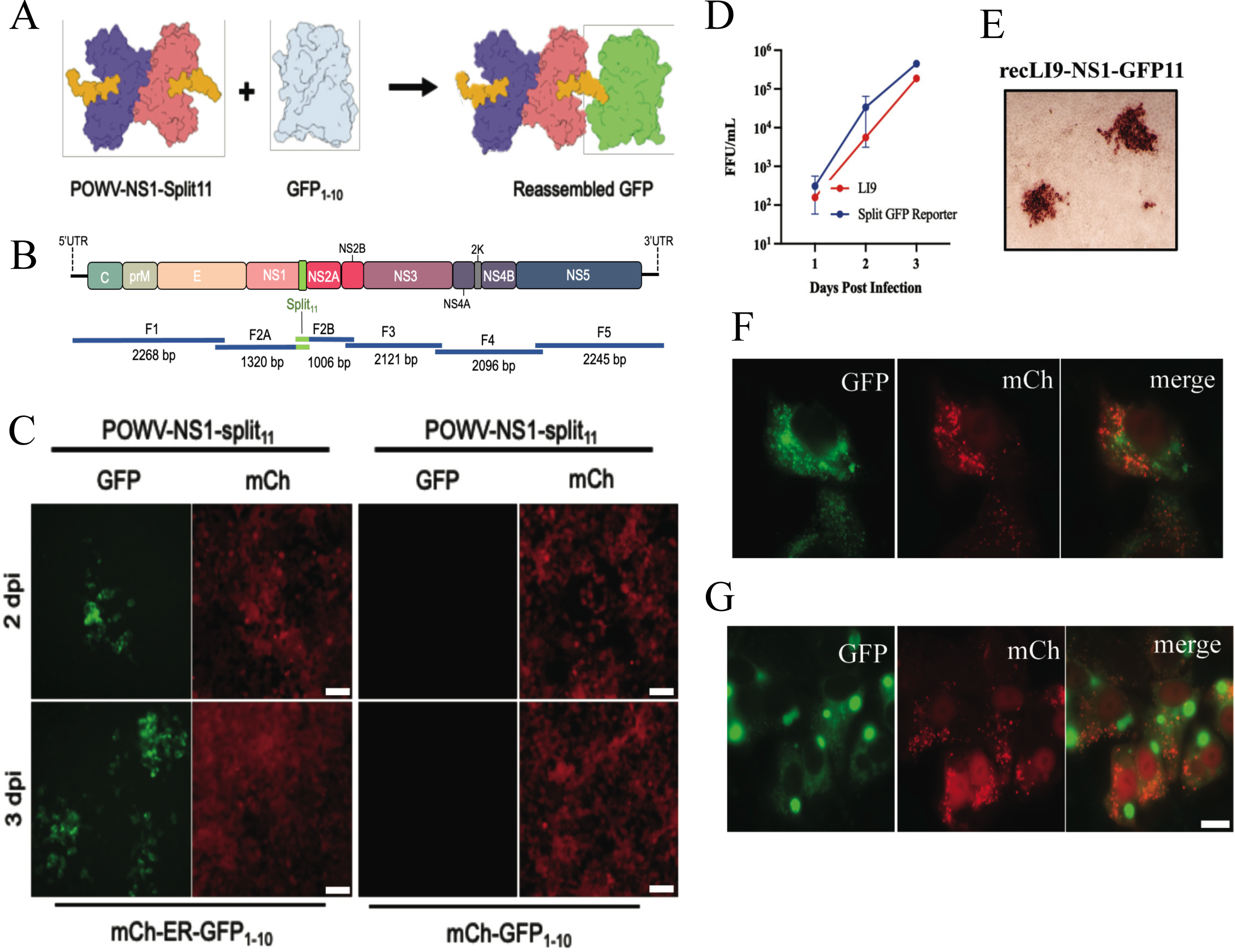
Generation of a Split GFP POWV Reporter Virus. **(A)** Schematic of the Split GFP fluorescence reporter system directed by the recLI9-NS1-GFP11 virus. Individual NS1 monomers (blue and pink), with a 16 residue GFP11 tag added to the NS1 C-terminus (yellow) with co-expressed, nonfluorescent, GFP_1-10_ (gray). The recLI9-NS1-GFP11 expression of NS1-GFP11 protein reconstitutes GFP fluorescence in cells co-expressing GFP_1-10_. **B)** CPER strategy for POWV-NS1-split11 generation. F2 was split into subfragments F2A and F2B with primer directed incorporation of GFP11 sequences. **(C)** Retrovirus transduced HEK293T constitutively expressing mCherry (cytoplasm) and either ER-localized ER-GFP_1-10_ (mCh-ER-GFP_1-10_) or cytoplasm localized GFP_1-10_ (mCh-GFP_1-10_), were infected with recLI9-NS1-GFP11. Live imaging captured 2-3 dpi shows foci of GFP fluorescence in LI9-NS1-GFP11 infected cells expressing ER translocated GFP_1-10_, but not cytoplasmically expressed GFP_1-10_. **(D)** Growth kinetics of WT LI9 (red) and recLI9-NS1-GFP11 (blue) 1-3 dpi in VeroE6 cells (MOI 1). **(E)** Immunostaining of LI9-NS1-GFP11 infected VeroE6 cell foci 3 dpi with anti-POWV HMAF. **(F and G)** Retrovirus transduced VeroE6 cells expressing mCherry-ER-GFP_1-10_ were LI9-NS1-GFP11 infected, and 6 dpi cells paraformaldehyde fixed. Representative images show perinuclear NS1-GFP fluorescence (F) and the discrete accumulation of NS1-GFP in large intracellular vesicles 6 dpi (G). Bars represent 10 µm.

In order to define mechanisms for attenuating POWVs, we developed a CPER reverse genetics system that authentically generates infectious recombinant POWVs. The CPER approach bypasses stability issues of bacterial FV clones and creates recombinant POWVs from amplified POWV genomic cDNAs and clones(^36, 59–62, 66^). The recLI9 POWV was CPER generated by circularizing 5-7 fragments of LI9 cDNAs with defined CMV promoter and HDVr directed 5’ and 3’ termini. CPER assembled LI9 DNA transfected into HEK or VeroE6 cells results in infectious recLI9 progeny that differ only by cell transfection efficiency. The authenticity of recLI9 viruses with WT LI9 was verified by sequencing, replication kinetics and phenotypically by focal cell-to-cell spread (Fig. 1,2).

Our newly developed POWV CPER reverse genetics system permits us to define determinants of POWV spread, neuroinvasion and neurovirulence, providing a mechanism for genetically attenuating POWVs and rationally designing live attenuated POWV vaccines. Reverse genetics has been accomplished for several FVs, initially using RNA transcription or plasmid approaches, and more recently by employing CPER systems(^36, 59–62, 66^). Reverse genetics has also been used to generate chimeric FVs with prM-Env specific neutralization determinants, and NS protein backgrounds derived from attenuated FVs or the Yellow Fever Virus (YFV) 17D vaccine (^31, 45, 51, 67^). Insertion of the POWV prM-Env in a YFV-17D background retained lethal neurovirulence with only 43% of inoculated mice surviving this virulent chimera(^50^).

Mutating FV Envelope and NS1 protein glycosylation sites have been associated with reduced replication and neuroinvasion(^23-25^, ^88-91^). However, how glycosylation alters FV virulence is likely dependent on both Env and NS1 functions that in combination determine FV assembly, spread and tissue tropism(^46^). FV NS1 proteins are reported to regulate viral replication, particle formation and functions of secreted extracellular NS1. Mutating residues 160 and 273 of the JEV NS1 protein regulates replication and particle formation, while mutating YFV or DENV NS1 N130 glycosylation sites reduces viral yield *in vitro* and neurovirulence following i.c. inoculation of mice(^34, 45^). In contrast, multiple mutations around NS1 glycosylation sites were required for WNV attenuation(^47, 92^).

POWV contains 2 novel NS1 glycosylation sites, not found in mosquito-borne FVs, and adds complex glycans to unique sites on the periphery of putative NS1 hexamers (Fig. S2). To define POWV NS1 glycosylation functions, we modified individual NS1 glycosylation sites (N85Q, N208Q, N224Q) and isolated infectious recLI9 NS1 mutant POWVs (Table S1). Each glycosylation site mutant spread focally cell-to-cell in VeroE6 cells like parental LI9, but with a 1-2 log decrease in NS1 mutant titers and reduced replication kinetics. A pronounced 2-3 day reduction in the timing of foci formation by the recPOWV-NS1-N224Q mutant coincides with reduced replication and reveals a severely attenuated *in vitro* phenotype. Although i.c. inoculation of mice with the NS1-N224Q glycosylation mutant was lethal, peripheral inoculation of the recLI9-NS1-N224 mutant resulted in 80% survival, elicited antibody responses to POWV Env proteins and a partially attenuated phenotype relative to WT LI9 infection. Further analysis is needed to determine if the NS1-N224Q glycosylation mutant is attenuated at peripheral or neuroinvasive levels of spread, and it remains to be determined how discrete POWV NS1 glycosylation sites regulate replication and delay POWV cell-to-cell spread. Irrespective of the mechanism, our findings rationalize including N224Q mutations in strategies for developing a live attenuated POWV vaccine.

Mutating WNV NS1 glycosylation sites (N130, N207) produced stable attenuated strains with NS1 accumulation in the ER and restricted NS1 secretion(^47, 92^). In DENV NS1, N207 mutants showed reduced dimer stability and secretion, with complex glycans attached to N130 and high mannose glycans added to N207(^31, 38^). The NS1 N85 glycosylation site was reported to be broadly required for high mannose or complex glycan processing, suggesting that N85Q mutations broadly restrict Golgi processing of N208/N224 glycans and alter NS1 secretion(^26^). Although all single POWV LI9 NS1 glycosylation mutants formed dimers and were secreted from cells, NS1 secretion was reduced in N224Q mutants, and increased in NS1 N85Q and N208Q mutants relative to WT LI9. It remains to be determined how the position of individual glycosylation sites (Fig 2B,S2) impact higher order NS1 glycan processing or hexamer formation, or constrain NS1 secretion and POWV release from cells.

Why some NS1s are highly secreted, while others remain highly cell associated, and how NS1 proteins regulate the assembly and secretion of FVs remains an enigma linked to viral spread and pathogenic mechanisms(^2, 28, 31, 40, 93^). POWV virions(^10^) and NS1 proteins remain highly cell associated with low levels of secreted NS1, and this novel NS1 attribute may have a role in directing the unique cell-to-cell spread phenotype of POWVs. Many functions ascribed to NS1 don’t reflect that NS1 proteins are cotranslationally translocated into ER compartments and unavailable for cytoplasmic interactions(^2, 28, 31^). Further, NS1 and Env proteins that are both present in secretory vesicles yet discretely released from cells, suggest intertwined roles for NS1 and Env glycosylation in virion assembly or secretion that have yet to be discovered(^23, 36, 40, 93^).

As an initial approach to study POWV NS1 translocation and secretion, we used CPER to generate a recLI9-NS1-GFP11 reporter virus for analysis of NS1 localization in cells. Fluorescence of the recPOWV NS1 split GFP11 protein was reconstituted in the ER of cells expressing ER-GFP1-10 protein. This demonstrates the ability of recLI9-NS1-GP11 to serve as a fluorescent reporter virus, and permits real-time analysis of POWV infection and NS1 functions in the lumen of the ER(^68, 70, 94^). Initial studies demonstrate the ER localization of NS1-GFP fluorescence and the novel accumulation of NS1-GFP in large intracellular vesicles in infected cells 6 dpi. These findings reveal NS1 lined vesicles and suggest a mechanism for NS1 retention that may restrict assembled POWV particle release(^10^) or contribute to a vesicular mechanism of POWV cell-to-cell spread. The nature of NS1 fluorescing vesicles (secretory, lipid droplet, autophagosome, convoluted membranes) remain to be defined(^28, 35, 38, 40, 46, 47, 92, 93^), but may explain how NS1 proteins are excluded from virions within secretory vesicles, yet determine FV spread in mammalian cells(^36, 95, 96^). Combining NS1 glycosylation mutants in a LI9-NS1-GFP reporter background may reveal roles for NS1 glycosylation sites in POWV secretion that impact attenuation. Further, the production of an infectious POWV NS1-GFP reporter virus provides a platform for high throughput screening of POWV replication inhibitors(^63, 65, 83, 94^).

Our findings provide a robust mechanism for generating recombinant POWV mutants and defining determinants of cell to cell spread, neuroinvasion and neurovirulence *in vitro* and *in vivo*. We generated a viable infectious POWV reporter virus that permits analysis of NS1 functions required for viral replication, maturation and neurovirulence, and reveal an NS1-N224Q glycosylation mutation that attenuates POWV replication *in vitro* and partially attenuates POWV in mice. Studies presented provide a mechanism for assessing attenuating POWV mutations and genetically designing live attenuated POWV vaccines.

## Methods

### Cells and Virus

Human embryonic kidney 293T cells (HEK293T) and Vero E6 cells were maintained in Dulbecco modified Eagle’s medium (DMEM; Gibco) with 8% fetal bovine serum (FBS), 10,000 U/mL of penicillin, and 10,000 µg/mL of streptomycin (Gibco, USA), and following viral infection, in media with 2% FBS. POWV strain LI9 (GenBank accession: MZ576219) was isolated from infected *Ixodes scapularis* ticks in VeroE6 cells and passaged 3-4 time in VeroE6 cells(^19, 97^). Work with infectious LI9 POWV and CPER-generated recombinant LI9 POWVs (recLI9) was performed in a certified BSL3 facility at Stony Brook University.

### Viral RNA Extraction and cDNA synthesis

RNA from POWV-infected VeroE6 cells was RLT extracted and RNeasy column (Qiagen) purified. cDNA synthesis was performed using the Transcriptor first-strand cDNA synthesis kit (Roche) following the manufacturer’s protocol and a primer complimentary to the 3’UTR (Table S2, oligo F5R)(^19, 97, 98^).

### PCR amplification of DNA fragments and CPER reactions

Five POWV-LI9 fragments were amplified from viral cDNA (Figure 1A,B) using high-fidelity Phusion polymerase (NEB) and corresponding paired primers (Table S2) that have a complementary 26-nucleotide overlap. An additional PCR fragment (UTR linker) was generated from plasmid pMiniT containing the CMV promoter, the first and the last 26 nts of the POWV-LI9 sequence, an HDVr, and SV40 polyadenylation site. POWV cDNA was amplified into 6 individual POWV DNA fragments: initial denaturation 98°C for 30s; 32 cycles of: 98°C for 20s, 60°C for 30s, and 72°C with 30s per kb; and a final extension at 72°C for 5 min. Resulting fragments were gel purified, Monarch kit (NEB) extracted and 0.1 pmol of each DNA fragment was CPER amplified in a 50 µL reaction containing 200 µM of dNTPs, 1x Phusion polymerase GC reaction buffer, and 1 µL Phusion polymerase. The following cycling conditions were used: initial denaturation 98°C for 30s; 12 cycles of: 98°C for 20s, 60°C for 30s, and 72°C for 10 min; and a final extension at 72°C for 10 min. CPER reactions lacking the UTR-Linker fragment were generated and used as negative controls.

For generating POWV-NS1_N85Q_, NS1_N208Q_, and NS1_N224Q_ mutant viruses, subfragments of fragment F2 (F2A, F2B) were generated by PCR from LI9 POWV cDNA, with primers containing AAC to CAA codon changes (Table S2) to generate NS1 proteins containing N85Q, N208Q or N224Q mutations. Subfragments F2A and F2B replaced F2 in CPER reactions as above. For generating the recLI9-NS1-Split_11_ reporter virus, subfragments of fragment F2 were similarly amplified from LI9 cDNA using primers containing Split_11_ sequences (Table S2) and used in CPER reactions as above.

### CPER Transfection and RecLI9 virus Rescue

CPER reactions were Monarch kit purified, eluted in sterile H_2_O and transfected into HEK293T cells seeded in 12-well plates using Lipofectamine 3000 reagent (Invitrogen)(^19, 97, 98^). Two days post-transfection, supernatants were harvested and amplified in VeroE6 cells to generate viral stocks and compared to transfection of control CPER reactions. Total RNA from VeroE6 cell infections with recLI9, recLI9-NS1_N85Q_,-NS1_N208Q_, and -NS1_N224Q_ mutant viruses, and recLI9-NS1-GFP11 reporter virus was purified using RNeasy kit (Qiagen). cDNA synthesis was performed as above using random hexamer primers (25°C for 10 min, 50°C for 60 min, and 90°C for 5 min) and sequenced as previously described(^19^) in the SBU Genomics facility.

### POWV LI9 Infection and Immunostaining

WT LI9 POWV, recLI9, recLI9-NS1_N85Q_, recLI9-recNS1_N208Q_, and recNS1_N224Q_ mutant viruses were adsorbed to **∼**60% confluent VeroE6 cells for 1 h. Following adsorption, monolayers were washed with PBS and grown in DMEM-2% FBS. Viral titers were determined by focus assay following serial dilution and infection of VeroE6 cells, and quantifying infected cells 24 hpi by immunostaining with anti-POWV hyperimmune mouse ascites fluid (HMAF; 1:5,000 [ATCC]), HRP-labeled anti-mouse IgG (1:3,000; KPL-074-1806), and AEC staining as previously described(^19^).

### Western blotting

VeroE6 cells were infected with WT LI9 or recLI9 mutant viruses (MOI, 1) and cells were harvested at 7 dpi in lysis buffer containing 1% NP-40 (150 mM NaCl, 50 mM Tris-Cl, 10% glycerol, 2 mM EDTA, 10 nM sodium fluoride, 2.5 mM sodium pyrophosphate, 2 mM sodium orthovanadate, 10 mM β-glycerophosphate) with 1x protease inhibitor cocktail (Sigma). Total protein levels were determined in a bicinchoninic acid assay (Thermo Scientific), and 20 µg of protein was resolved by SDS– 10% polyacrylamide gel electrophoresis. Proteins were transferred to nitrocellulose, blocked in 5% bovine serum albumin, and incubated with the indicated antibodies. Antibodies used were mouse anti-POWV NS1 (M837; Native Antigen Company) (1:5,000), anti-POWV HMAF sera (ATCC) (1:1,000), and anti-GAPDH (G9545; Sigma-Aldrich) (1:5,000)(^19, 97, 98^). Protein was detected using HRP-conjugated anti-mouse secondary antibody (Amersham) and Luminata Forte Western HRP substrate (Millipore).

### Glycosidase Analysis

VeroE6 cells were infected with LI9, recLI9-NS1_N85Q_, recLI9-NS1_N208Q_, and recLI9-NS1_N224Q_ mutant viruses (MOI, 1) and harvested 7 dpi. Cells were washed with PBS, lysed in 1% NP-40 buffer as above and 20 µg of lysate or 50 µL of supernatants were subjected to Endo H (P0702S NEB) or PNGase F (P0709S NEB) digestion following the manufacturer’s protocols. Samples were resolved by SDS–10% polyacrylamide gel electrophoresis, Western blotted(^19, 97, 98^) and viral NS1 protein was detected using mouse anti-POWV NS1 (M837; Native Antigen Company) as above.

### Retrovirus GFP_1-10_ Vectors

Retrovirus vectors were generated by transfecting HEK293T or VeroE6 cells using polyethylenimine (PEI) transfection using a DNA/PEI ratio of 1:3(^99^). HEK293T cells (3.8 × 10^6^) were preincubated with 25 µM chloroquine diphosphate for 5 h and transfected with pQCXIP-mCh-2A-ER-GFP_1-10_ or pQCXIP-mCh-2A-GFP_1-10_ plasmids with packaging plasmids pMMLV-CMV-GagPol and pLP/VSVG using in a 3:2:1 ratio. After 18 h, media was replaced, and the viral supernatants were harvested 72 hpt and filtered through a 0.45-µm PVDF filter.

### Live Cell POWV Fluorescence

Packaged retroviruses, mCh-2A-ER-GFP_1-10_ or mCh-2A-GFP_1-10_, were used to transduce ∼50% confluent monolayers of VeroE6 or HEK293T cells to constitutively express mCherry-ER-GFP_1-10_ or mCherry-GFP_1-10_. After 2 days, cells were selected using puromycin (5 or 0.5 µg/mL, respectively, for VeroE6 and HEK293 cells) for 2 days. Puromycin selected cells were infected with recLI9-NS1-GFP11 (MOI 5) in microslides chambers (IbITreat, Germany) and live-cell fluorescence was monitored using a Nikon Axiovert microscope in the BSL3. Additionally. infected VeroE6 were fixed for 1 hour in 1% paraformaldehyde, washed 3 times with PBS, and fluorescence was measured using EVOS M5000 microscope with a 60x oil immersion objective. Control for split-GFP system specificity were performed by PEI co-transfection of pQCXIP-mTag-ER-split11 or pQCXIP-mTag-split11 with pQCXIP-mCh-2A-ER-GFP_1-10_ into ∼50% confluent HEK293T. Fluorescence was observed 2 days post-transfection using Olympus IX51 microscope and Olympus DP71 camera (Fig S3).

### LI9 POWV NS1 Molecular Modeling

The 3D model of dimeric LI9 NS1 protein was obtained using multimer prediction in AlphaFold2 using ColabFold within ChimeraX 1.5 software(^100^). Five models were created with the top-scoring model selected after relaxation. The POWV LI9 NS1 hexamer model was generated using cryoEM structures from DENV2 soluble NS1 hexamers (PDB 7WUV) scaffolding with the model relaxed using Rosetta Relax(^101, 102^).

### In Vivo LI9 POWV Infection Experiments

Anesthetized murine C57BL/6 pups (<2-week-old, N=3-5) were intracranially infected with 2 × 10^2^ FFU POWV or buffer-only control in a volume of 2 μl. Infected mice were monitored twice daily for signs of disease and daily for weight loss. Peripheral Inoculation: C57BL/6 mice (male, 41∼51-week-old, N=4-10, Jackson Laboratory) were anesthetized via intraperitoneal injection with 100 mg·ml^−1^ of ketamine and 20 mg·ml^−1^ of xylazine per kilogram of body weight. Animals were infected via subcutaneous footpad injection with 2 × 10^3^ FFU POWV or buffer-only control in a volume of 20 μl. Weights and neurovirulent sequalae were monitored daily as indicated.

### Biosafety and Biosecurity

Animal research was performed in accordance with institutional guidelines following experimental protocol review, approval, and supervision by the Institutional Biosafety Committee and the Institutional Animal Care and Use Committee at Stony Brook University. Animals were managed by the Division of Laboratory Animal Resources (Stony Brook University), which is accredited by the American Association for Accreditation of Laboratory Animal Care and the Department of Health and Human Services. Animals were maintained in accordance with the applicable portions of the Animal Welfare Act and the DHHS ‘Guide for the Care and Use of Laboratory Animals.’ Veterinary care was under the direction of full-time resident veterinarians boarded by the American College of Laboratory Animal Medicine. Experiments with infectious POWV were performed in an ABSL3 facility at SBU.

## Acknowledgements

We thank Jorge Benach and Stella Tsirka for supportive discussions on tick-borne diseases and neuropathogenesis and manuscript feedback and Smruti Mishra and Luke Helminiak for animal monitoring and sample retrieval. This work was supported by funding from a DOD TBDRP Idea Development Award W81XWH2210702, National Institutes of Health grants: NIAID R01AI12901005, R21AI13173902, R21AI15237201, RO1AI027044, T32AI007539, an IRACDA Post-Doctoral Award and a Stony Brook University Seed Grant. The funders had no role in study design, data collection and interpretation or the decision to submit the work for publication. We declare no conflict of interest.

## Supplemental Figures

**Figure S1.** VeroE6 cells were infected with WT LI9 POWV, recLI9-NS1_N85Q_, recLI9-NS1_N208Q_, and recLI9-NS1_N224Q_ mutant viruses (MOI, 1) and 7 dpi cell RNAs were extracted, and cDNA synthesized using a 3’UTR-reverse primer. Mutants were sequenced and sequencing chromatograms of mutated glycosylation sites are shown versus WT LI9 to validate mutant construction.

**Figure S2.** The POWV NS1 hexamer model was generated by using the cryoEM structure of DENV2 soluble NS1 hexamers (PDB 7WUV) as scaffolding and then relaxed using Rosetta Relax(^101, 102^). NS1 domains were pseudocolored in blue (β-roll), pink (wing), and yellow (β-ladder). Asparagine N85, N208, and N224 are colored red within the structure.

**Figure S3.** HEK293T cells were PEI co-transfected with cytoplasmic or ER localized GFP11 expression constructs (pQCXIP-mTag-ER-split11 or pQCXIP-mTag-split11) and plasmid pQCXIP-mCh-2A-ER-GFP_1-10_ that expresses ER-translocated GFP_1-10_. Live cell microscopy visualized 2 dpt demonstrates the specific reconstitution of GFP only when GFP11 is ER localized.

---

**Table S1.** CPER POWV Reverse Genetics Viral Rescue.

| Recombinant POWVs | Viral Rescue |
| --- | --- |
| N85Q | + |
| N208Q | + |
| N224Q | + |
| N85Q/N208Q | - |
| N85Q/N224Q | - |
| N208Q/N224Q | - |
| NS1-Split11-C-terminal | + |
| NS1-Split11-297 | - |

**Table S2.**
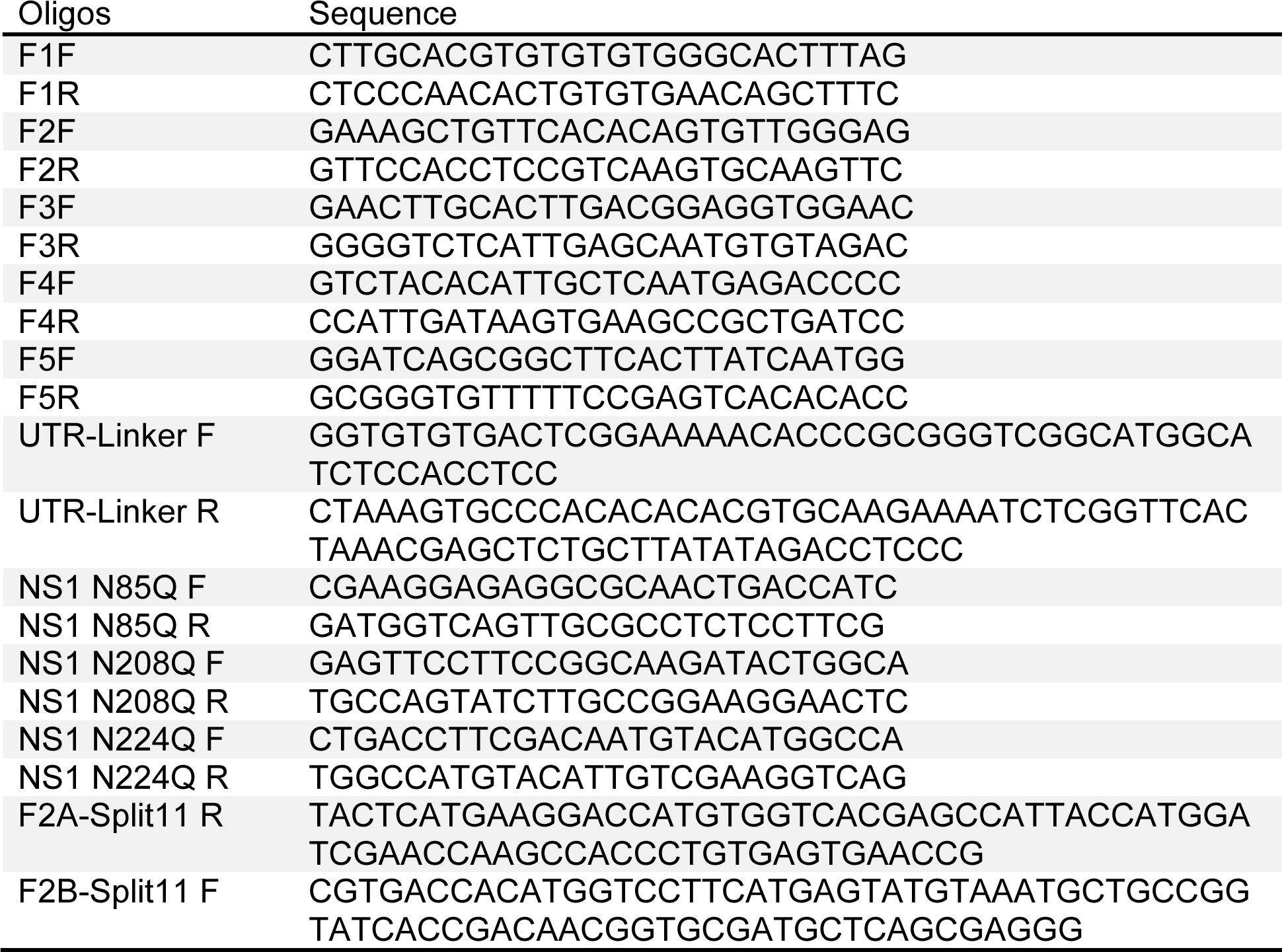
Oligonucleotides.

