## Supplemental Fig 1-3 for "Establishment of a CPER Reverse Genetics System for Powassan Virus Defines Attenuating NS1 Glycosylation Sites and an Infectious NS1-GFP11 Reporter Virus"

### Supplemental Figure 1

POWV-NS1$_{N85Q}$

POWV-NS1$_{N208Q}$

POWV-NS1$_{N224Q}$

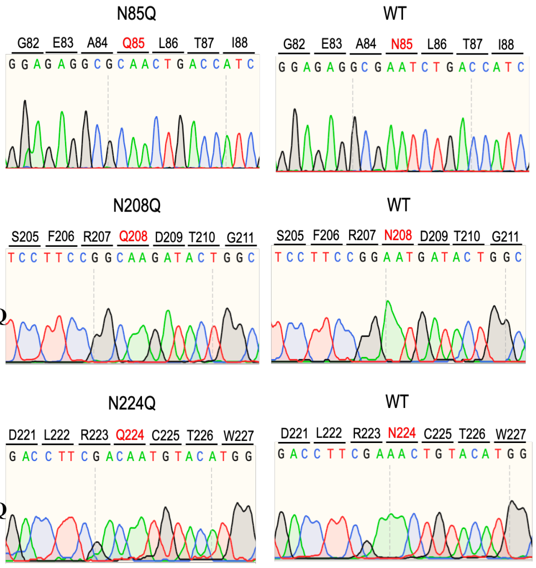


### Supplemental Figure 2

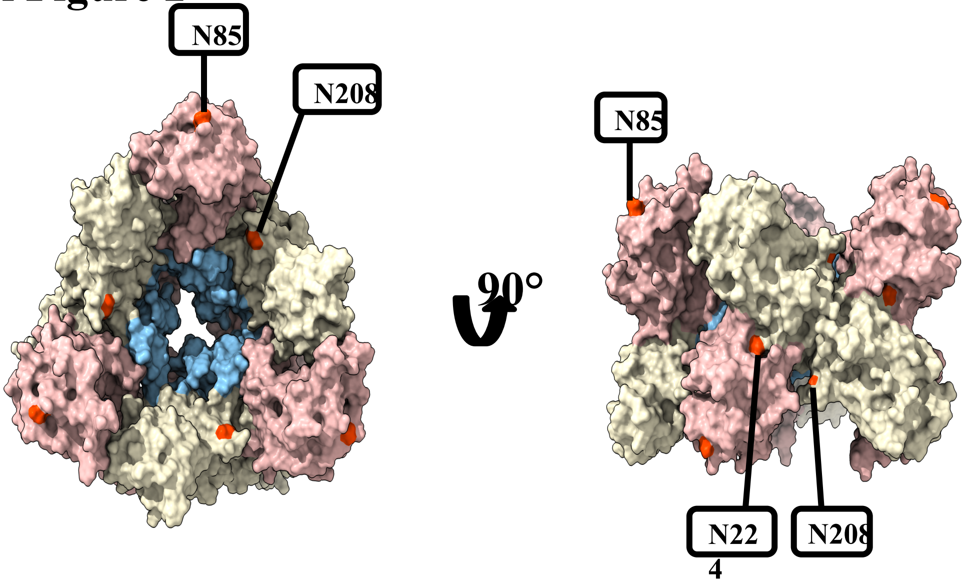


### Supplemental Figure 3

mTag-ER-split$_{11}$ | mTag-split$_{11}$

GFP | mCh | GFP | mCh

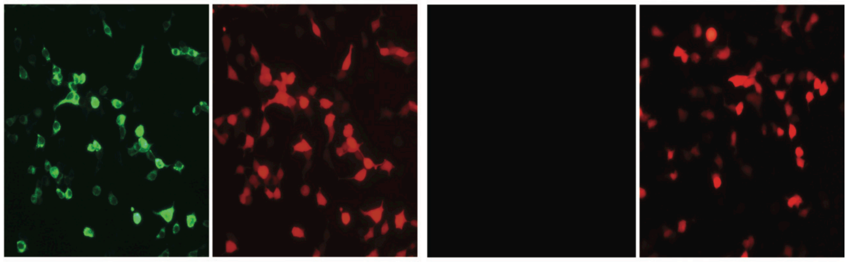

mCh-ER-GFP$_{1-10}$-transduced cells
